# Integrated Use of Compost and Nitrogen Fertilizer and their Effects on the Yield and Yield Components of Wheat: A Pot Experiment

**DOI:** 10.1101/2020.10.07.329524

**Authors:** Bayu Dume Gari, Gezahegn Berecha, Melkamu Mamuye, Obsu Hirko Diriba, Amsalu Nebiyu, Abebe Nigussie, Aleš Hanč, Pavel Švehla

## Abstract

There is a research gap related to the combined effects of compost produced from coffee husks and inorganic nitrogen fertilizer (urea). The objective of this study was to evaluate the yield and yield components of wheat (*Triticum aestivum L.*) under the integrated application of compost and nitrogen fertilizer (urea). A pot experiment was conducted in a lath house to determine the effects of the integrated use of compost produced from coffee husks and nitrogen fertilizer (urea) on the yield and yield components of wheat. The experiment consisted of nine treatments: T1, control (untreated); T2, 5 t ha^−1^ (8.12 g pot^−1^) compost; T3, 10 t ha^−1^ (16.24 g pot^−1^) compost; T4, 0 t ha^−1^ compost + 50 kg ha^−1^ nitrogen fertilizer (NF) (0.09 g pot^−1^); T5, 5 t ha^−1^ compost + 50 kg ha^−1^ NF; T6, 10 t ha^−1^ compost + 50 kg ha^−1^ NF; T7, 0 t ha^−1^ compost + 100 kg ha^−1^ (0.18 g pot^−1^) NF; T8, 5 t ha^−1^ compost + 100 kg ha^−1^ NF; and T9, 10 t ha^−1^ compost + 100 kg ha^−1^ NF. The treatments were arranged in a completely randomized design (CRD) with three replications. The compost was prepared from coffee husks and applied in wet conditions. The findings showed that the addition of compost had little effect on wheat yield and yield components in the absence of nitrogen fertilizer (urea). However, the application of the highest amount of nitrogen fertilizer (urea), which is equivalent to the recommended field rate (100 kg ha^−1^) (equivalent to 0.18 g pot^−1^), and compost (5 t ha^−1^) (equivalent to 8.12 g pot^−1^) led to a significant (P≤0.05) increase in grain yield. Under this treatment, the grain yield was 26 g pot^−1^ (equivalent to 14.741 t ha^−1^) which is a 66.29% increase compared with the control (8.9 g pot^−1^ (4.969 t ha^−1^)); in the treatment in which only the recommended amount of nitrogen fertilizer was used (21.98 g pot^−1^ (12.273 t ha^−1^)) grain yield increased by 16.74%. Spike length and dry matter yield also significantly (P≤0.05) increased with the application of integrated compost and nitrogen fertilizer (urea). The results of this experiment revealed that compost-based soil management strategies can enhance wheat production, thereby contributing positively to the viability and benefits of agricultural production systems. However, nutrient-compost interactions should receive special attention due to the great variability in the properties of compost, which may depend on the type of organic materials used.

## INTRODUCTION

In many Sub-Saharan African countries, increasing wheat production to support the higher demands of growing populations is still a challenge [1, 2]. Ethiopia is one of the largest wheat producers in Sub-Saharan Africa and has a favorable climate for growing wheat. However, over the past decade, there has been a consistent deficit in wheat production. The fragmented nature of land holdings and limited use of agricultural inputs, particularly fertilizers, contribute to low levels of wheat productivity in Ethiopia [3]. This country does not produce its own fertilizer supply, and farmers apply a common fertilizer, diammonium phosphate (DAP), and urea at a recommended rate of 100 kg ha^−1^ each, regardless of soil type. Several studies have reported that inorganic fertilizer application has resulted in increased crop productivity worldwide. However, the supplementation of crop areas using only two sources of inorganic fertilizers (DAP and urea) for many years and the complete removal of crop residues from the production land has led to severe nutrient depletion in Ethiopia. On the other hand, the use of compost alone, as a substitute for inorganic fertilizer, is not enough to maintain the present levels of crop productivity of high-yielding varieties [4]. Therefore, integrated nutrient management, in which both compost and inorganic fertilizers are used simultaneously, is the most effective method, particularly in developing countries where mineral fertilizers are cost-prohibitive. Few previous studies [5, 6, 7] showed that wheat productivity could be enhanced through combinations of organic and inorganic fertilizers. Some trials have been conducted on the agronomic response of wheat to organic (i.e., farmyard and green manure) and mineral fertilizer integrations. However, there have not been any investigations on the combined effects of compost produced from coffee husks and inorganic fertilizer (urea) on the yield and yield components of wheat. In Ethiopia, 192,000 metric tons of coffee husks are produced and disposed of per year, and the study area (Jimma) is well-known for coffee production; in this area, 134,400 metric tons of coffee husks per year are simply disposed of into the environment [8]. Thus, it is very important to use coffee husks for agricultural input by composting and integrating them with inorganic fertilizers. Therefore, the objective of this study was to evaluate the yield and yield components of wheat (*Triticum aestivum L.*) under integrated application of compost produced from coffee husks and nitrogen fertilizer (urea).

## MATERIALS AND METHODS

### Description of the study area

This study was conducted at Jimma University College of Agriculture and Veterinary Medicine, Southwestern Ethiopia, in a lath house pot experiment. The study area is located at a latitude of 7°33’N and longitude of 36°57’E. The altitude ranges from 1760 to 1920 m above sea level. The mean annual maximum and minimum temperatures are 26.8 and 11.4°C, respectively, and the maximum and minimum relative humidity values are 91.4 and 39.92%, respectively. The mean annual rainfall is 1500 mm, and soil is mainly dominated by Nitisols [9].

### Soil sampling and compost preparation

Composite samples were collected from the top 0-30 cm soil layer. The collected soil samples were air-dried, crushed using a mortar and pestle and then passed through a 2 mm square-mesh sieve. Compost derived from coffee husks was prepared in Jimma University College of Agriculture and Veterinary Medicine (JUCAVM) using the pit composting method.

### Soil and compost analysis

pH and electrical conductivity (EC) were measured in distilled water at a ratio of 1:5 (w:v) [10]. The OC content was determined by the Walkley-Black method, percentage of OM was obtained by multiplying the percent of soil OC by a factor of 1.724, following the assumption that OM is composed of 58% carbon and total N was analyzed using the Kjeldahl method [11]. The available phosphorus (P) was determined by the Mehlich 3 method [12]. The CEC and exchangeable bases (Ca, Mg, K and Na) were determined after extracting the samples with 1M NH_4_OAc at a pH of 7. The Fe content was determined with an atomic absorption spectrometer (AAS) after extraction with DTPA solution. The Ca and Mg contents in the extracts were analyzed using AAS, while Na and K contents were analyzed by flame photometer [11]. The exchangeable acidity of the soil was determined by the titration method after a 1 M KCl solution (pH 7) was used to leach the exchangeable H^+^ and Al^3+^ from the soil sample [13]. The particle size distribution (texture) of the soil sample was determined by the Boycouos hydrometric method [11]. The soil bulk density was determined by the undisturbed core sampling method after drying the soil samples in an oven at 105°C to a constant weight.

### Pot experiment set up

A pot experiment was carried out at a controlled lath house. Plastic pots (top and bottom diameters of 33 cm and 20 cm, respectively) 30 cm in height were filled with air-dried soil. The experiment consisted of the following nine treatments: T1, control; T2, 5 t ha^−1^ (8.12 g pot^−1^) compost; T3, 10 t ha^−1^ (16.24 g pot^−1^) compost; T4, 0 t ha^−1^ compost + 50 kg ha^−1^ (0.09 g pot^−1^) nitrogen fertilizer (NF); T5, 5 t ha^−1^ compost + 50 kg ha^−1^ NF; T6, 10 t ha^−1^ compost + 50 kg ha^−1^ NF; T7, 0 t ha^−1^ compost + 100 kg ha^−1^ (0.18 g pot^−1^) NF; T8, 5 t ha^−1^ compost + 100 kg ha^−^ NF; and T9, 10 t ha^−1^ compost + 100 kg ha^−1^ NF. The treatments were arranged in a completely randomized design (CRD) with three replications. Ten wheat (*Triticum aestivum L.*) seeds were sown in each pot and the seedlings were thinned to keep five plants per pot after germination. After emergence, four plants were maintained in each pot until harvesting. The plants were regularly watered with distilled water to maintain field capacity moisture content until the end of maturity. Plant height was recorded just before harvesting. The aboveground parts of the plants were harvested at the soil level from each pot. The fresh shoots were dried separately at 70°C for 72 hrs.

### Postharvest soil and nutrient uptake analysis

The NO_3_-N in the plant tissues was determined using 2% acetic acid [14]. The plant K content was determined by a flame photometer after wet digestion with sulfuric acid. The plant P content was determined photometrically with the molybdenum blue method. At the end of the experiments (after crop harvest), soil samples were collected from each pot, air-dried and sieved (<2 mm), and selected chemical properties were analyzed by following standard procedures [11]. The pH and EC of the soil were determined in a water suspension at a 1:5 soil:liquid ratio (w:v). The total carbon content was determined by the Walkley-Black method, and the total N content was analyzed using the Kjeldahl method [11].

### Statistical analysis

The data were subjected to statistical analyses. One-way analysis of variance (ANOVA) was performed to compare variations in the soil properties and plant growth characteristics for each treatment. For all the analyses, treatment means were separated using the least significant difference (LSD), and treatment effects were declared significant at the 5% probability level (P≤0.05). All analyses were performed using SAS version 9.3.

## RESULTS AND DISCUSSION

### Soil and compost properties

The physicochemical properties of the experimental soil and compost are shown in Table 1. The soil had lower available P and total P and higher exchangeable Fe and Al than the compost. The high pH values of the compost may be due to basic cations, such as Ca, Mg, Na, and K, which were present in the compost materials. The CEC of the compost was also high, which may be due to the high negative charge potential of surface functional groups. The OC concentrations were high in the compost. The compost contained high levels of both macro- and micronutrient elements. The compost had especially high amounts of basic cations (Ca, Mg, K, and Na) and total P. These results are supported by the findings of [15], who reported that compost derived from coffee husks contained high levels of organic matter and micro- and macronutrients.

**Table 1.** Selected physicochemical properties of the experimental soils and compost

| Parameter | Experimental soil<br>(mean±SD) | Compost<br>(mean±SD) |
| --- | --- | --- |
| Bulk density (gm cm<sup>3</sup>) | 1.04 ± 0.01 | - |
| pH-H<sub>2</sub>O (1:5) | 5.2 ± 0.02 | 7.6±0.03 |
| Exch. acidity (cmol kg<sup>-1</sup>) | 4.9± 0.1 | - |
| EC (dS/cm) (1:5) | 0.02 ± 0.00 | 4.32 ± 0.21 |
| Exch. Ca (cmol kg<sup>-1</sup>) | 8.08 ± 1.32 | 58.35 ± 0.52 |
| Exch. Mg (cmol kg<sup>-1</sup>) | 1.20 ± 0.2 | 6.45 ± 0.02 |
| Exch. K (cmol kg<sup>-1</sup>) | 0.8 ± 0.02 | 1.98 ± 0.21 |
| Exch. Na (cmol kg<sup>-1</sup>) | 0.02 ± 0.00 | 3.05 ± 0.21 |
| Exch. Fe (cmol kg<sup>-1</sup>) | 35.54± 1.12 | - |
| Exch. Al (cmol kg<sup>-1</sup>) | 795.00± 0.23 | - |
| CEC (cmol kg<sup>-1</sup>) | 24.36 ± 1.7 | 68.12 ± 0.21 |
| Organic carbon (%) | 3.97 ± 0.23 | 30.80 ± 6.12 |
| Organic matter (%) | 6.85 ± 0.39 | 53.10 ± 1.50 |
| Total nitrogen (%) | 0.34 ± 0.02 | 1.62 ± 0.62 |
| Total P (mg/kg) | 19.2± 0.14 | 130.01±2.25 |
| Available P (mg/kg) | 4.52 ± 0.09 | 14.45 ± 1.23 |

### Wheat growth characteristics and yield components

Compared to the control, the addition of compost increased wheat grain production in the absence of nitrogen fertilizer (NF). These increases were clearly lower than those caused by the use of NF without the addition of compost. These results are in agreement with those of several authors [16,17], who found little response of crop yield and nutrient status to the application of compost alone, likely due to the carbon-rich nature of compost. Therefore, most of the previous studies have shown that the beneficial effects of the addition of compost on crop production are most evident when compost is combined with NF. The detected increases in nutrient efficiency after amending soil with compost have been mainly related to greater nutrient retention, minimal nutrient losses, improvements in soil properties (such as increases in WHC and decreases in soil compaction, leading to the immobilization of contaminants or the mobilization of nutrients), and the enhancement of soil biological properties, such as the development of more favorable root environments and microbial activities favoring nutrient availability. In this study, a significant compost*NF interaction was observed for wheat grain production (Table 3). Thus, the highest wheat grain production was obtained by combining 5 t ha^−1^ compost and 100 kg ha^−1^ NF. These values represented an increase relative to the values of the control soil and the control soil with the addition of the highest amount of NF. These results demonstrated that the addition of 5 t ha^−1^ compost increased grain yield relative to the use of only NF (at the recommended rate). In this experiment, nutrient losses were not observed, since irrigation was controlled to avoid leaching, and the compost adsorption capacity might have limited nitrogen availability in the short term, especially after NF addition. However, under field conditions, the fact that the addition of compost can limit nutrient losses from leaching may favor an increase in the availability of nutrients in the soil in the long term. The substantial increase in the total C and N contents in the compost treatments suggests that compost may be useful for building C and N contents in soils, particularly those with inherently low levels of C and N.

The results indicate that the growth parameters of wheat are influenced by different integrations of coffee husk compost and nitrogen fertilizer (urea) (Table 2). The highest shoot length in the heading stage was recorded in the treatment in which nitrogen fertilizer (urea) was applied at the recommended rate (100 kg ha^−1^) together with 5 t ha^−1^ compost, and the lowest shoot length was recorded in the treatments with no nitrogen fertilizer applications. This suggests that wheat growth might be slowed if wheat cultivation is not supplemented with mineral fertilizers, regardless of the application of compost.

**Table 2.**
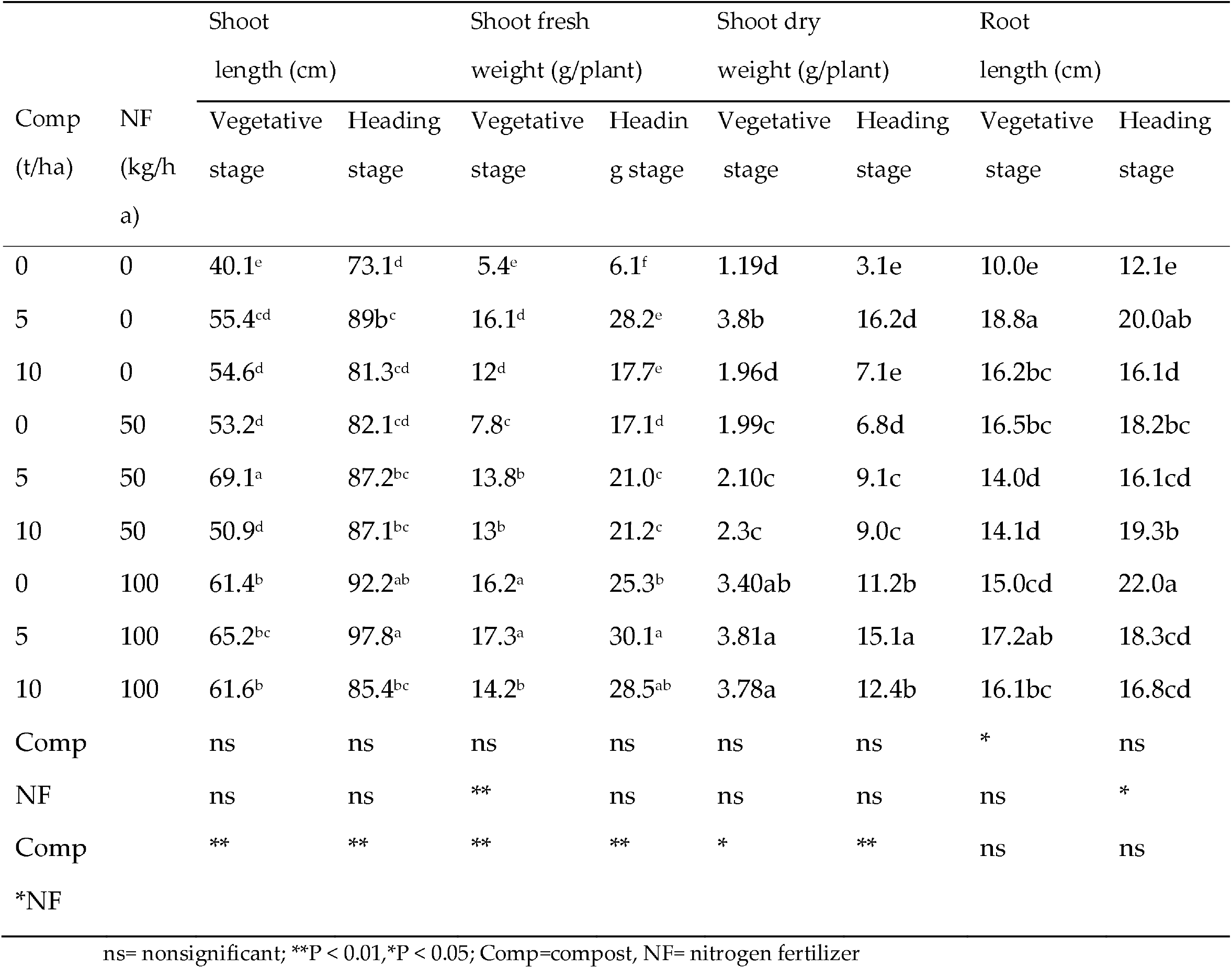
Effect of compost applied alone or mixed with NF on the shoot and root characteristics of wheat

**Table 3.**
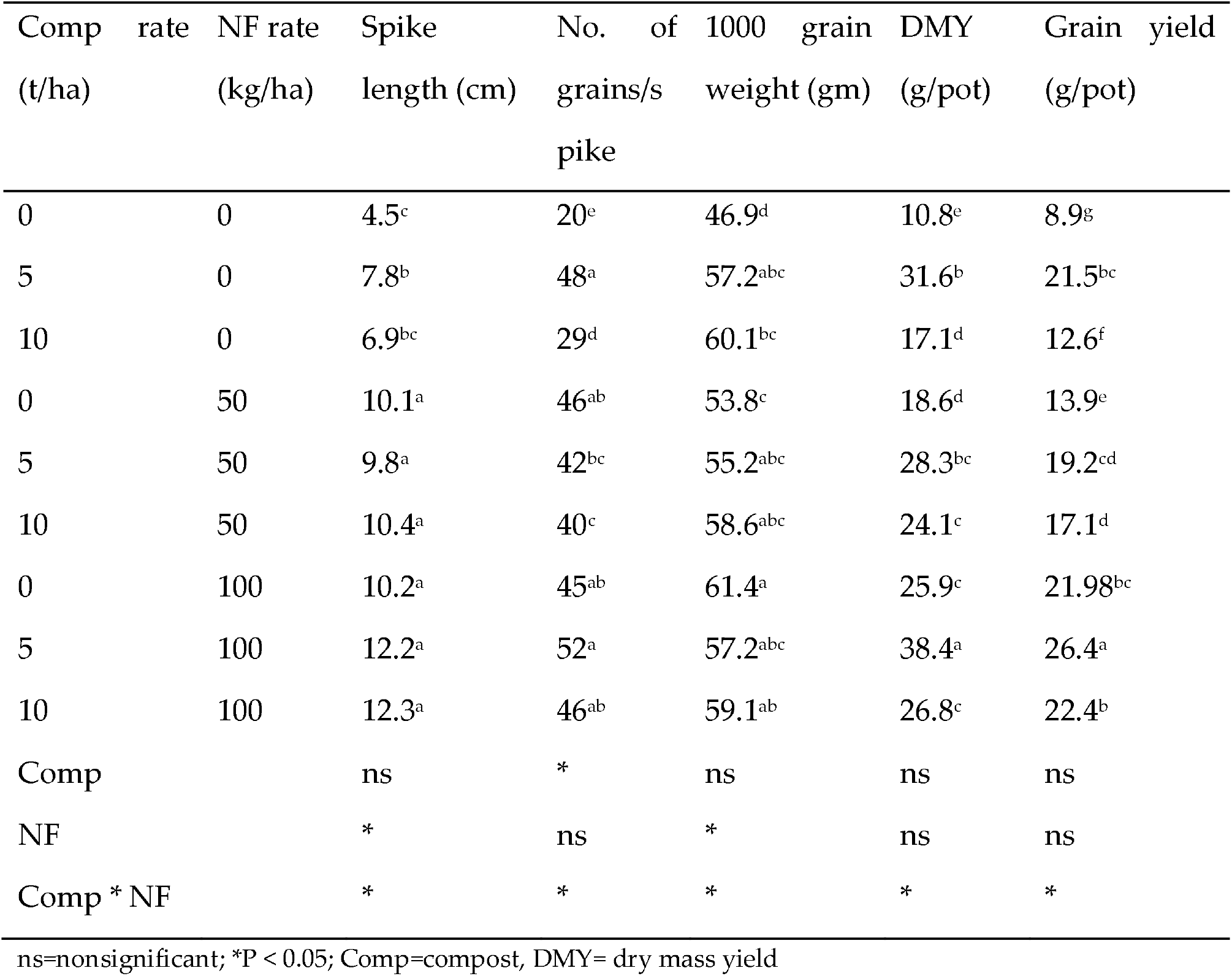
Effect of compost alone and mixed with NF on the yield and yield characteristics of wheat

A greater spike length (12.3 cm) was observed when wheat was cultivated under 100 kg ha^−1^ nitrogen fertilizer combined with 10 t ha^−1^ compost; this treatment was not significantly different from the treatment that received 50 kg ha^−1^ nitrogen fertilizer and 5 t ha^−1^ compost (12.2 cm). Generally, combinations with 50-100 kg ha^−1^ nitrogen fertilizer and 5-10 t ha^−1^ compost lead to significantly greater spike lengths and numbers of grains per spike. However, a smaller spike length was recorded in the treatment with no nitrogen fertilizer and compost. Similarly, the dry matter yield was significantly influenced by the different combinations of nutrient sources and rates. The highest DMY (38.4 g pot^−1^) was observed in the treatment with the combined application of 5 t ha^−1^ compost and 100 kg ha^−1^ nitrogen fertilizer, and the lowest DMY (10.8 g plot^−1^) was recorded from the control treatment (no compost or nitrogen fertilizer application). Fertilizer combinations significantly influenced other yield and yield components, such as thousand seed weight, grain yield and dry matter yield (Table 3). Thousand seed weight showed significant variation across treatments. The highest thousand seed weight (61.4 gm) was recorded when 100 kg ha^−1^ nitrogen fertilizer alone was applied. Similar results were reported by different researchers, e.g., Kaur, Sarwar, Rasool and Brar [18– 21]. Lower thousand seed weights were observed in treatments to which compost alone was applied (no nitrogen fertilizer), indicating that unless it is integrated with inorganic fertilizers, the use of compost alone may not fully satisfy crop nutrient demand, especially in the year of application [22]. Compared to wheat cultivated with no fertilizer application (control), grain yield was more than three times higher when wheat was cultivated under 100 kg ha^−1^ nitrogen fertilizer and 5 t ha^−1^ compost. This might be because the soil from the study area had poor soil fertility.

### Plant nutrient concentration

The application of compost significantly increased the shoot concentrations of K and P and significantly decreased the concentrations of N, Ca, Mg, Cu, Mn and Zn (Table 4). The sulfur concentration in the shoot was not affected by compost application; however, there was a significant (P ≤ 0.05) increase in its concentration with NF application. The interactive effects of NF and compost treatments were nonsignificant in relation to the concentrations of all the nutrient elements (Table 4).

**Table 4.**
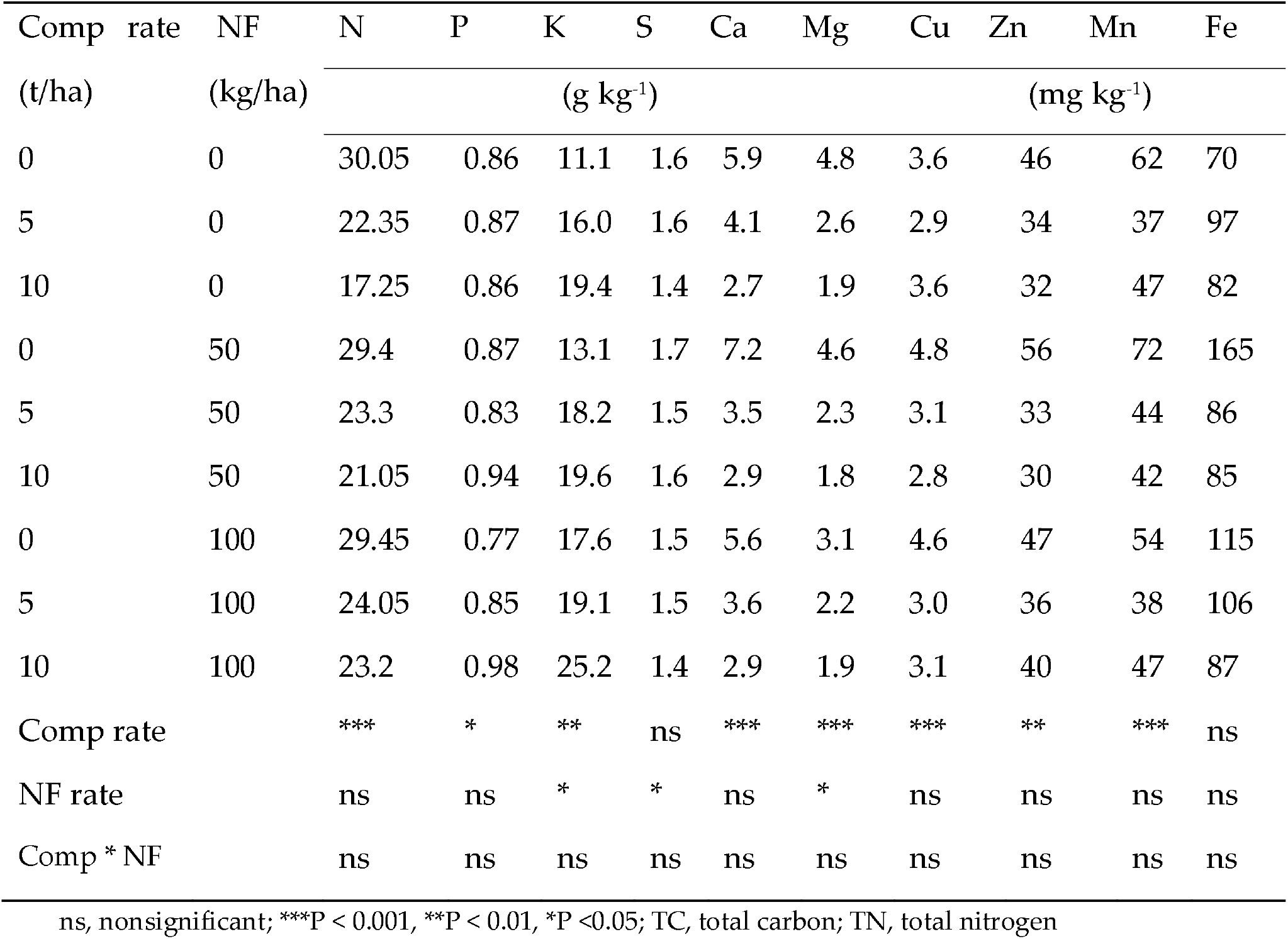
Concentrations of macro- and micronutrients in wheat shoots grown with different application rates of compost and NF

### Plant nutrient uptake

Shoot uptake data and associated statistical analyses for various nutrients are presented in Table 4. The shoot uptake of all the nutrients except Ca and Mg increased significantly with the application of compost (Table 4). The uptake of Ca and Mg remained similar or increased slightly in response to the compost treatments. Fertilizer application had a significantly positive effect on the uptake of nutrients, except for Ca, Mg, Mn, and Fe, for which the effect was not significant (Table 4). The plant uptake of almost all the macro- and micronutrients increased with increasing compost application, and the decrease in nutrient concentration in the shoots was most directly related to the increased shoot dry matter yield of the plants (dilution effect) in the compost-treated soil. The decreased concentration of some of the nutrients, however, could be due to their adsorption onto the Fe and Al oxides present in the acidic soil. Iron and Al oxides are known to have high adsorption capacity for phosphate and metallic cations such as Cu, Zn, Mn, Mg and Ca.

### Changes in postharvest soil properties

At the end of the experiments (after crop harvest), composite soil samples were collected and analyzed. The compost had a large influence on the soil properties, which can explain its effects on plant growth and grain production. The increasing application of compost significantly increased the pH, EC, TC and TN in the soil (Table 5). The total C and N contents increased significantly with the compost treatments, although the increase in the soil C was substantially greater (Table 5), and the fertilizer treatment significantly increased the total N content. The application of compost significantly increased soil total C relative to that in the control. Interactions between compost and soil may enhance soil C storage via the sorption of organic matter to compost [18]. Coffee husk compost offers great potential as a soil amendment, as it is a potential source of nutrients. The increased soil pH due to compost addition might have increased the adsorption capacity of metallic cations. The compost had a great influence on the soil properties, which can explain its effects on plant growth and grain production. The results of this study highlighted the significant improvement in the soil amended with compost alone, while the growth and yield of the crops supplemented with both compost and NF were higher than that of the crops supplemented with only the highest amount of N, displaying the value of mixed treatments.

**Table 5:** Effects of compost and fertilizer treatments on pH, EC, total C and total N measured after plant harvest

| Compost<br>rate t/ha) | Fertilizer<br>rate kg/ha) | pH-H<sub>2</sub>O | EC (dS/cm) | TC (g kg<sup>-1</sup>) | TN (g kg<sup>-1</sup>) |
| --- | --- | --- | --- | --- | --- |
| 0 | 0 | 5.20 | 0.164 | 316.12 | 34.25 |
| 5 | 0 | 5.46 | 0.336 | 319.20 | 34.15 |
| 10 | 0 | 5.96 | 0.626 | 436.40 | 35.30 |
| 0 | 50 | 5.11 | 0.188 | 316.21 | 34.23 |
| 5 | 50 | 5.45 | 0.409 | 439.15 | 35.26 |
| 10 | 50 | 5.83 | 0.996 | 468.10 | 38.60 |
| 0 | 100 | 5.24 | 0.250 | 429.30 | 35.17 |
| 5 | 100 | 5.42 | 0.389 | 430.22 | 35.28 |
| 10 | 100 | 6.15 | 1.052 | 462.14 | 38.70 |
| Comp rate |  | *** | *** | *** | *** |
| NF rate |  | * | *** | ** | * |
| Comp x NF |  | ns | *** | ns | ns |

ns, nonsignificant; \*\*\*P < 0.001, \*\*P < 0.01, \*P < 0.05; Comp, compost; NF, nitrogen fertilizer

While the lath house experiment is a good starting point for evaluating plant responses under ideal conditions, the true agronomic potential of coffee husk compost needs to be assessed by measuring crop responses to this compost under field conditions. Therefore, the use of integrated compost and nitrogen fertilizer is a feasible approach to overcome soil fertility constraints. Additionally, the integrated use of nitrogen fertilizers and compost may improve the efficiency of mineral fertilizer.

## Acknowledgments

Financial support for this work was provided by the Capacity building for scaling up of evidence-based best practices in agricultural production in Ethiopia (CASCAPE) project. Jimma University College of Agriculture and Veterinary Medicine, Jimma, Ethiopia, provided funding for this research.

## Data Availability Statement

All relevant data are provided within the paper.

## Author Contributions

Conceptualization: BDG GB MM.

Formal analysis: BDG MM.

Funding acquisition: GB.

Investigation: BDG GB MM OHD AN AN AH PS.

Methodology: BDG MM.

Project administration: GB.

Resources: GB.

Writing original draft: BDG.

Writing, reviewing & editing: BDG GB MM OHD AN AN AH PS.

